# Rapid Phenotypic Antibiotic Susceptibility Determination Direct From Spiked Urine Samples Shows Strong Correlation with Gold Standard for Urinary Tract Infection

**DOI:** 10.64898/2026.09.21.753156

**Authors:** Megan Roegner, Chelsey Ahmadi, Scotlynn Lyon, Zac McGee, Krishnan Chittur, Paula Millirons

## Abstract

GeneCapture is developing a novel, point-of-care, rapid AST technique that, when fully automated, can accomplish AST within 2 hours directly from a raw sample, representing a significant advancement in diagnostic capabilities. For this study, a panel of uropathogen isolates was collected from clinical specimens positive for urinary tract infection from various laboratories in the southern US. A selection of these isolates were then spiked into fresh clean-catch urine samples to produce mock samples covering a variety of antibiotic responses. Testing was performed on 20 strains of *Escherichia coli*, the cause of over 80% of UTIs, and 6 strains each of *Klebsiella pneumoniae* and *Proteus mirabilis*, the next most common causes. Each strain was tested against the susceptible, intermediate and resistant (S-I-R) breakpoints of up to 5 appropriate antibiotics including Amoxicillin-Clavulanate, Cefpodoxime, Ciprofloxacin, Nitrofurantoin and Trimethoprim-Sulfamethoxazole. The new method uses a phenotypic assessment of bacterial growth, comparing responses from bacteria treated with antibiotics at S-I-R breakpoint concentrations to untreated controls as the metric for determining susceptibility. Exposure to the drugs for only 90 minutes gave a Categorical Agreement of 98% to gold standard culture results across the resulting 247 bacteria/antibiotic pairs. Testing on separate days of nine of these strains in triplicate gave a reproducibility of 97%. The sequential steps of this novel AST - sample processing, growth, labeling, and counting - can readily be performed on an automated cartridge that is currently under development. The presented work, which took place in a laboratory setting, demonstrates the efficacy of the assay.

**Importance:** The rapid identification of pathogenic bacteria and determination of their antibiotic susceptibility directly from a patient sample remains a critical challenge in modern healthcare. Urinary tract infections (UTIs) account for a large proportion of global antibiotic use, and are increasingly resistant to former first-line treatments. Uncomplicated UTI (uUTI) is still generally treated empirically at the point of care, which leads to frequent follow-up visits. GeneCapture’s novel AST method relies on a phenotypic assessment of bacterial growth, as genotypic methods are not well suited for all bacteria. The speed of the assay is gained through using fresh samples, a novel rapid fluorescent in-situ hybridization technique, and accurate growth determination. The ease of use and affordability of the automated system should allow adoption at both clinical and point of care locations to allow more accurate and rapid antibiotic treatment.

## Introduction

The rapid increase in antimicrobial resistance (AMR) is a major threat to global health.^1^ Accurate diagnosis of infections combined with appropriate antibiotic therapies is vital for combating AMR.^2^ To determine an appropriate antibiotic for an individual’s infection, the susceptibility of that specific infectious organism must be determined. Current gold standard methods for antimicrobial susceptibility testing (AST) include culturing of bacterial colonies over the course of 16-36 hours, followed by biochemical or mass spectrometry-based identification (e.g., MALDI-TOF), and phenotypic AST, which can take up to an additional 24 hours.^3,4,5^ Lab-based phenotypic AST involves the culture of bacteria in the presence of doubling concentrations of antibiotics, which are reported as minimum inhibitory concentrations (MICs). These can be interpreted as susceptible, intermediate, or resistant (S, I or R) by laboratorians. The wait for AST results delays treatment or allows ineffective antibiotics to be prescribed, both of which can lead to worsening infections.

Uncomplicated UTI (uUTI) is one of the most common infections requiring medical intervention and antibiotic use, with greater than 10 million visits per year in the US alone.^6^ Treatment at the point-of-care is often symptom-based even though symptoms alone (with or without dipstick results) do not adequately determine the presence of a true infection.^7,8^ Although most UTI pathogens remain susceptible to at least one of the standard treatments, the prevalence of resistance increased by 7% per year in uropathogenic *Escherichia coli* from 2011 to 2019 with a 3% yearly increase in the presence of three or more resistance mechanisms in a single isolate.^9^ This trend continues to escalate. Multidrug resistant *Klebsiella pneumoniae* in out-patient UTIs are spreading across US communities from multiple diverse lineages.^10^

Studies have shown that up to 20% of antibiotics prescribed empirically for outpatient UTI were ineffective (R or I), and that these patients are 60% more likely to need follow-up care (ranging from different drug prescriptions to hospitalization) than those who received a drug that was indeed effective.^11,12^ Additionally, patients prescribed unnecessary or inappropriate antibiotics contribute to rising AMR, both in the patients through unnecessary exposure, and in the community at large. This delay also increases the risk to the patient of complications such as pyelonephritis, urosepsis, bacteremia, or chronic UTI.^9,13^ These negative consequences drive up healthcare costs for patients, insurers, and medical providers.^14,15^

In response, there has been a recent rise in more rapid diagnostic techniques, aiming to reduce the time between initial empiric treatment, and actual targeted therapy.^16^ Many of these technologies use molecular methods to identify AMR from the presence of known resistance genes, but the presence of these genes does not always determine phenotypic susceptibility, particularly in gram negative organisms, which cause over 90% of uUTIs.^15^ Thus, phenotypic AST remains the gold standard in determining antibiotic susceptibility.^17,18^ Most novel rapid tests currently depend on using a positive blood culture tube, which itself requires 8-48 hours of incubation. Only one POC direct from sample AST is currently available, and only in Europe where it is not required to identify the pathogen before reporting AST results.^19^

GeneCapture, Inc. is responding to the need for rapid point-of-care (POC) diagnostics to speed the accurate and targeted treatment of infections by developing a rapid phenotypic POC AST assay, named CAPTURE AST. This assay is currently being evaluated in a laboratory setting, with development of a POC cartridge and instrument underway. CAPTURE AST tests breakpoint concentrations of a panel of 5 antibiotics and reports the traditional S-I-R results or simply S or Not Susceptible (NS) for each drug against the three most common uropathogens, *E. coli, K. pneumoniae* and *Proteus mirabilis*. This work demonstrates CAPTURE AST’s ability to report accurate phenotypic susceptibility results directly from a spiked urine sample in under two hours.

## Methods Samples

### Strains

Isolated uropathogens have been collected by GeneCapture as plated colonies from a variety of commercial testing sites and preserved as glycerol stocks. All species identifications from the source were verified in house using standard microbiology techniques. Gold standard susceptibility data was confirmed for each isolate using in-house broth microdilution or Kirby-Bauer disk diffusion. For this study, the selection of strains was chosen to represent a variety of antibiotic responses. As *E. coli* is the most predominant species causing uUTI, 20 strains of *E. coli* were included, as well as 6 strains each of *K. pneumoniae* and *P. mirabilis*.

### Broth-cultured bacteria

Glycerol-preserved stocks of bacteria were streaked for isolation on blood agar plates (Hardy Diagnostics, Santa Maria, CA) and incubated overnight at 35-37ºC. Isolated colonies were picked and placed in 10 mL of Tryptic Soy Broth supplemented with Yeast Extract (TSY) (Sigma-Aldrich, St. Louis, MO) in a shaking incubator rotating at 300 RPM at 35-37ºC overnight. On the day of testing, 250 µL of overnight culture was transferred to 25 mL of Cation-Adjusted Mueller Hinton Broth (CaMHB, ThermoScientific, Waltham, MA) and grown for 30 minutes in a shaking incubator at 35-37ºC. Unlabeled bacteria were quantified by diluting 1:100 in sterile 1x phosphate buffered saline (PBS, Sigma-Aldrich, St. Louis, MO) and counted on an Accuri C6+ Flow Cytometer (BD, Franklin Lakes, NJ). Raw cell counts were converted to the concentration of colony forming units in the subculture (CFU/mL), and subcultures were then diluted in additional CaMHB to a final assay concentration of approximately 500,000 CFU/mL (ranging from 300,000 to 800,000 CFU/mL). This concentration was used as the starting concentration for the AST assay.

### Spiked Urine Samples

To create the spiked urine samples, broth-cultured uropathogens were added to fresh clean catch urine collected from local volunteers under an IRB-approved protocol. Five mL of the spiked urine sample was then passed through a 0.2 µm filtration mock-up using a vacuum apparatus at approximately 5 PSI. The bacteria were backflushed off the filter using a syringe with 2 mL of CaMHB, quantified using flow cytometry as in the previous section, then diluted to a final assay concentration of approximately 500,000 CFU/mL. This procedure mirrors what will be performed on the automated cartridge.

### Internal validation testing

Prepared spiked samples were checked for purity by plating the initial growth control (T=0 Untreated) on blood agar and incubating in ambient air overnight at 35-37ºC.

The remaining CAMHB suspended samples for each condition tested (no-drug and with each drug) were allowed to incubate in ambient air overnight at 35-37ºC. This broth macro-dilution testing was used as a final check for the determined susceptibility of the gathered test data.

### AST Assay

#### Antibiotics

All antibiotics were acquired in powdered form from commercial retailers, and stock solutions were made following CLSI guidelines.^20^ Further dilutions were made for each antibiotic to 20x the highest concentration listed in **Table 1** and aliquots of these concentrations were stored at -20ºC except Amoxicillin/clavulanate which was stored at - 80ºC. Fresh aliquots were thawed for use prior to each test. Final dilutions into growth media were performed at the setup of each test. Aliquots were also tested weekly on QC organisms to validate effectiveness during storage.

**Table 1.** The three most common uropathogens and the drugs and concentrations tested for each breakpoint. N/A indicates a drug that is not appropriate for that species. Note: CLSI does not recognize an I breakpoint for SXT.

|  | Antibiotic concentrations in µg/mL for the S, I and R breakpoints |  |  |  |  |
| --- | --- | --- | --- | --- | --- |
| Organisms | Amoxicillin/<br>clavulanate (AMC) | Cefpodoxime<br>(CPD) | Ciprofloxacin<br>(CIP) | Nitrofurantoin<br>(NIT) | Trimethoprim/Sulfa-<br>methoxazole (SXT) |
| *E. coli*<br>(20 strains) | 8/4, 16/8, 32/16 | 2, 4, 8 | 0.25, 0.5, 1 | 32, 64, 128 | 2/38, 4/76 |
| *K. pneumoniae*<br>(6 strains) | N/A | 2, 4, 8 | 0.25, 0.5, 1 | 32, 64, 128 | 2/38, 4/76 |
| *P. mirabilis*<br>(6 strains) | N/A | 2, 4, 8 | 0.25, 0.5, 1 | N/A | 2/38, 4/76 |

#### Workflow

Testing was performed at S, I, and R breakpoints of each drug (**Table 1**) with a bacterial inoculum of about 500,000 CFU/mL. Each test included determining the concentration of bacteria in an untreated starting point control (T=0), an untreated growth control (T=90), and in each antibiotic condition (T=90, AUG, S breakpoint; etc.).

#### Growth

Each setup was made in a 1 mL volume for more accurate manual pipetting, then a 20 µL portion was loaded into a 96-well PCR plate. A volume of 100 µL of a proprietary labeling solution was immediately added to the T=0 control. Each T=0 control was heated to 55ºC for 15 minutes, cooled in the dark at room temperature for 5 minutes, then diluted 10-fold into three separate aliquots of 1x PBS in a round bottom 96-well plate. Each of these was counted on the commercial flow cytometer for 1 minute at a medium flow rate. Counts of the three aliquots were averaged for a final count. The remaining samples were placed in a 35-37ºC incubator for the growth time (90 minutes), then followed the same labeling and counting process as the T=0 sample.

#### Labeling

The CAPTURE AST assay uses a unique fluorescent in-situ hybridization (FISH) step with probes linked to fluorescent dyes that label specific target bacteria. FISH probe design was informed using Geneious software (Dotmatics.com) for sequence alignment and analysis of conserved regions in the 16S and 23S ribosomal RNA sequences. Using an in-house developed buffer (patent application in preparation), the FISH labeling was performed as a one-step process in 15 minutes at the end of the growth phase. The probes used include a published universal bacterial probe, EUB338 (ACT CCT ACG GGA GGC AGC AG), and custom-designed probes specific to each AST panel organism: *E. coli, P. mirabilis*, and *K. pneumoniae*.

#### Analysis

The CAPTURE AST assay captures the fluorescent signals from the labeled bacteria to generate enumeration counts from each cell population. These data are analyzed using proprietary algorithms to determine if that organism is S-I-R or S-NS for each antibiotic. These algorithms compare cell populations in each antibiotic treatment group to the control growth wells for that run.

## Results

### Spiked Urine Bacterial Testing

To mimic clinical samples, testing was completed using uropathogens isolated in the southern US and spiked into freshly collected clean catch urine samples. The antibiotic panel was selected for appropriateness for the uUTI condition. Whether analyzed using the traditional S-I-R calls (**Tables 2** and **4**) or S-NS (**Tables 3** and **5**), these data show an overall 98% categorical agreement to reference methods.

**Table 2.** S-I-R agreement of the spiked urine dataset as a ratio of correct to total runs with indicated errors.

| SIR Agreement | AMC | CPD | CIP | NIT | SXT |
| --- | --- | --- | --- | --- | --- |
| *E. coli* | 34/34 | 35/35 | 34/35, 1 INV | 35/36, 1 MIN | 34/35, 1 VMJ |
| *K. pneumoniae* | N/A | 10/11, 1 MIN | 8/9, 1 MAJ | 10/11, 1 MIN | 9/10, 1 MAJ |
| *P. mirabilis* | N/A | 10/10 | 10/10 | N/A | 10/10 |

**Table 3.** S-NS agreement of the spiked urine dataset as a ratio of correct to total runs with indicated errors.

| S-NS Agreement | AMC | CPD | CIP | NIT | SXT |
| --- | --- | --- | --- | --- | --- |
| *E. coli* | 34/34 | 35/35 | 34/35, 1 INV | 36/36 | 34/35, 1 VMJ |
| *K. pneumoniae* | N/A | 10/11, 1 MAJ | 8/9, 1 MAJ | 11/11 | 9/10, 1 MAJ |
| *P. mirabilis* | N/A | 10/10 | 10/10 | N/A | 10/10 |

**Tables 2** and **3** show the ratios of correct to total runs and the types of errors, if any, by organism and drug. MIN represents a minor error (calling an S or R organism as I or vice versa), MAJ represents major errors (false resistance); and VMJ represents very major errors (false susceptibility). Minor errors (MIN) only apply to the S-I-R calling method. Invalid (INV) calls are due to inconsistencies in growth patterns between the three concentrations or other issues with a test.

When analyzed by drug, the data from these 247 drug-organism pairs are represented in **Tables 4** and **5**. The combined results from either analysis method meets the FDA’s required data metrics of ≥90% Categorical Agreement (CA), <10% INV results, <3% MAJ, and <1.5% VMJ. Due to the small numbers in certain categories, individual metrics appear higher than acceptable (e.g., CPD, MAJ errors), but with sufficient replicates, these are expected to more closely match the total values.

**Table 4.** Analysis of Table 2 data by drug meets FDA standards using the traditional S-I-R approach.

| DRUG | AGREE | MIN | MAJ | VMJ | INVALID | CA |
| --- | --- | --- | --- | --- | --- | --- |
| AMC | 100.00% | 0.00% | 0.00% | 0.00% | 0.00% | 100.00% |
| CPD | 98.15% | 0.00% | 3.23% | 0.00% | 0.00% | 98.15% |
| CIP | 96.49% | 1.75% | 0.00% | 0.00% | 1.75% | 98.21% |
| NIT | 95.74% | 4.26% | 0.00% | 0.00% | 0.00% | 95.74% |
| SXT | 96.36% | 0.00% | 3.57% | 3.70% | 0.00% | 96.36% |
| **TOTAL** | **97.17%** | **1.21%** | **1.38%** | **1.06%** | **0.40%** | **97.56%** |

**Table 5.** Analysis of Table 3 data by drug meets FDA standards using the simpler S-NS approach.

| DRUG | AGREE | MAJ | VMJ | INVALID | CA |
| --- | --- | --- | --- | --- | --- |
| AMC | 100.00% | 0.00% | 0.00% | 0.00% | 100.00% |
| CPD | 98.15% | 3.23% | 0.00% | 0.00% | 98.15% |
| CIP | 98.25% | 0.00% | 0.00% | 1.75% | 100.00% |
| NIT | 97.87% | 2.44% | 0.00% | 0.00% | 97.87% |
| SXT | 96.36% | 3.57% | 3.70% | 0.00% | 96.36% |
| **TOTAL** | **97.98%** | **2.07%** | **0.98%** | **0.40%** | **98.37%** |

Additionally, triplicate tests were performed on separate days with separate inocula on two to five isolates of each species as a measure of reproducibility. The five *E. coli* strains tested in triplicate showed 100% concordance, as did a pair of *P. mirabilis* strains. Of the two *K. pneumoniae* strains tested in triplicate, the first strain gave one MIN error in its first run, and the second strain gave two MAJ errors in its first run. These errors did not repeat. Overall, the reproducibility seen in this study was 97%.

## Discussion

As AMR rises, infections that once were amenable to empiric treatment are becoming more complex,^21^ with uUTIs significantly impacted by this trend. Antimicrobial stewardship requires evidence of pathogenic load in urine and phenotypic AST to inform correct treatment. Without phenotypic susceptibility, empiric treatment contributes to treatment failure, recurring infections, hospitalizations, and sepsis-related mortality^1.5^ Rapid tests that inform physicians of the causative agent and appropriate antibiotic treatment at the patient’s initial visit have the potential to reduce AMR and these complications.

The CAPTURE AST assay is designed to take a fresh urine sample and affordably determine its susceptibility in under two hours. To validate the assay, testing of cultured organisms was completed on the three most common uUTI pathogens: *E. coli, K. pneumoniae*, and *P. mirabilis*. Uropathogens, collected from local laboratories, were spiked into fresh urine to generate mock samples that showed an impressive 98% Categorical Agreement and 97% reproducibility over this study.

While this study focused on uUTIs, several GeneCapture studies have been completed to show CAPTURE AST’s relevance to other sample types. Early work with a less stringent protocol using common wound pathogens demonstrated an 88% CA and showed the ability of the assay to assess samples containing multiple pathogens, common in infected wounds and complicated UTIs (Supplemental Figure 1). Antifungal susceptibility tests (AFST) of multiple *Candida albicans* and *C. glabrata* (currently known as *Nakaseomyces glabratus*) have also been carried out and showed 97% CA in only 5 hours, much shorter than the current gold standard method of broth microdilution, which can take anywhere from 2 to 4 days under optimal conditions.^22^

All validation testing to date indicates that 90 minutes gives sufficient bacterial growth to report susceptibility to antibiotics on the CAPTURE AST panel from a fresh urine sample. This study demonstrates that the CAPTURE AST system can produce highly accurate and meaningful results directly from a urine sample in a fraction of the time for traditional AST (2-4 days) or even lab-based automated AST systems (6+hours after sample delivery). Rapid, point-of-care AST that includes pathogen ID has the potential to drastically improve patient outcomes.^23,24^ By utilizing fresh patient samples and precisely counting the targeted organisms in a sample, early impacts on bacterial growth from the antibiotic panel allow a short test to be accurate. CAPTURE AST uses a proprietary system for probe design and for labeling the bacteria after growth, as well as a proprietary machine-learning approach for analyzing the data to make accurate calls. The next step, testing discarded, de-identified fresh clinical samples suspected of uUTI, has begun and is producing similar results.

This study represents a novel, rapid time-to-result phenotypic AST and is achieved by the accuracy of the innovative cell labeling and data analysis methods, making the method compatible with an on-cartridge system. The effort to fully automate the process is underway.

## Acknowledgements

We thank Mary Beth Minyard, MSCLS, M(ASCP), for the contribution of many of the uropathogen isolates used in this work.

